# Experimental characterisation of *de novo* proteins and their unevolved random-sequence counterparts

**DOI:** 10.1101/2022.01.14.476368

**Authors:** Brennen Heames, Filip Buchel, Margaux Aubel, Vyacheslav Tretyachenko, Andreas Lange, Erich Bornberg-Bauer, Klara Hlouchova

## Abstract

*De novo* gene emergence provides a route for new proteins to be formed from previously non-coding DNA. Proteins born in this way are considered random sequences, and typically assumed to lack defined structure. While it remains unclear how likely a *de novo* protein is to assume a soluble and stable tertiary structure, intersecting evidence from random-sequence and *de novo*-designed proteins suggests that native-like biophysical properties are abundant in sequence space. Taking putative *de novo* proteins identified in human and fly, we experimentally characterise a library of these sequences to assess their solubility and structure propensity. We compare this library to a set of synthetic random proteins with no evolutionary history. Bioinformatic prediction suggests that *de novo* proteins may have remarkably similar distributions of biophysical properties to unevolved random sequences of a given length and amino acid composition. However, upon expression *in vitro, de novo* proteins exhibit higher solubility which is further induced by the DnaK chaperone system. We suggest that while synthetic random sequences are a useful proxy for *de novo* proteins in terms of structure propensity, *de novo* proteins may be better integrated in the cellular system given their higher solubility.

## Background

*De novo* genes, formed from previously non-coding DNA, have in recent years been confirmed as a ubiquitous feature of eukaryotic genomes, and are likely to represent an important source of new protein-coding evolutionary material (1–3). Translation of DNA that has not been under selection for its protein-coding capacity means that protein-coding *de novo* genes lie at the edge of yet-to-be-explored ‘dark protein space’ (4). Despite the unevolved nature of *de novo*-emerged proteins (here referred to as *de novo* proteins), many have been shown to play important functional roles. Examples include the putative *de novo* proteins Goddard, Atlas, and Saturn, which play essential roles in fly (5–7); three mouse-specific *de novo* proteins with diverse cellular roles (8); the yeast protein Bsc4, required for DNA-damage repair (9); and notothenioid antifreeze glycoprotein (10). Of these, Goddard and Bsc4 have been structurally characterised and found to have clearly maintained structural elements. However, both proteins appear to contain segments with high intrinsic disorder (ID). Bungard et al. (9) concluded that Bsc4 is best described as having a molten globule structure, suggesting that it may lack the defined folding funnel typical of many stable native folds.

Despite these examples, the structural properties of *de novo* proteins remain experimentally understudied. Computational prediction of the ID and aggregation propensity of *de novo* proteins has sparked hypotheses regarding the evolutionary pressure acting on newly-emerged proteins (11–14). Foremost is the suggestion that avoidance of aggregation is a critical selection pressure acting on novel proteins (15). Selection against aggregation would also explain why many studies identify higher ID in *de novo* proteins (given the fundamental link between amino acid hydropathy and ID). More complete answers to these questions will come from experimental characterisation: this should reveal the true distribution of aggregation propensity/ID in newly emerged protein sequences. Ultimately, systematic experimental characterisation of novel sequences should reveal if novel proteins have the capacity to form folded structures and how frequently this occurs.

*De novo* proteins have sometimes been approximated to ‘random’ sequences based on the lack of selection upon their emergence. However, *de novo* proteins emerge from existing genomes that are already known to carry different sequence and compositional biases, e.g. in GC-content (16). Diverse areas of research have shown that compositional biases can significantly impact properties such as translation efficiency, aggregation propensity and even specific attributes of ID (15, 17, 18). The extent to which *de novo* and random sequences can be regarded as proxies therefore remains unclear. Moreover, random sequences represent true occupants of dark protein space, whose properties themselves are heavily understudied. This region of sequence space has typically been assumed to contain non-functional and disordered proteins, proteins which are likely to be toxic and degraded if expressed in cells.

Nevertheless, many recent studies have identified both structure and function in random proteins. Structure itself appears to be abundant in protein sequence space. Secondary structure occurrence has been reported to be remarkably close to that of biological proteins. In addition, 20-40% of random sequence space has been observed to be resistant to proteolysis, likely due to tertiary structure formation (19–23). Furthermore, we were recently able to demonstrate that while structured random proteins are hard to express *in vivo* due to their higher aggregation propensity, random proteins with greater ID are readily tolerated by *E. coli* (22). Simultaneously, at least some protein folds appear to be relatively evolvable from random sequences. Hayashi et al. (24) were able to evolve an arbitrary random sequence to replace the D2 domain of an essential bacteriophage protein. Function through binding may be the most likely role that an unevolved protein could attain. For example, ATP-binders have been selected from pools of random proteins (25). Random and partially randomised peptides have also been shown to have functional effects when expressed in vivo (26–29). Finally, a smaller number of studies have evolved catalytic activity from randomised sequences, including esterase, barnase, and RNA-ligase activity; the presence of which is itself an indicator of structured catalytic centres (30–32). Altogether, while the above-listed studies suggest that both random and *de novo* proteins have non-zero structural and functional potential, their mutual relevance remains unclear.

Here, we set out to go further than previous studies by analysing the structural potential of *de novo* proteins. In doing so, we bring two strands of research together and experimentally characterise sets of *i)* 1800 putative *de novo* proteins identified in human and fly genomes and ii) 1800 synthetically-generated random sequences. While earlier studies were entirely computational, or experimentally characterised single proteins, we quantify the properties of putative *de novo* proteins and compare them to ‘true’ random sequences i.e. unevolved and synthetically-generated. We investigate two fundamental properties – solubility and structure content – using techniques previously unapplied to bulk analysis of *de novo* proteins.

We find that *de novo* proteins appear broadly similar to random sequences when length and amino acid frequencies are held constant. Consistent with computational prediction, the set of 1800 putative *de novo* proteins we study had similar overall protease resistance to the set of synthetic random sequences. This indicates that, at least given the amino acid composition of the *de novo* sequences chosen, random sequences have similar structural potential. However, we also find that *de novo* proteins are (moderately) more soluble at this composition and structure level. This is indicative of some selective pressure having acted over the course of their real – albeit short – evolutionary histories.

## Results

### A library-based approach enables high throughput investigation of *de novo* proteins

In this study we combine computational and experimental characterisation of two libraries: i) a set of 1800 putative *de novo* proteins identified in human or fly, and ii) a set of 1800 synthetic random sequences with no evolutionary history. Libraries were synthesised as an oligonucleotide pool, limiting proteins to 66 residues or less. A lower bound of 44 residues was chosen given the diminishing likelihood of domain-like structures for very short proteins. With these constraints, 1800 sequences were selected from published sets of novel protein (Fig. 1a). Fly sequences (n=176) are estimated to have emerged from previously non-coding intronic or intergenic regions <50 Mya, and all are annotated as protein-coding genes in *Drosophila melanogaster* (13). Human sequences (n=1624) are unannotated intronic or intergenic ORFs with *Homo sapiens-specific* expression (i.e. born <6.7 Mya) (12). We refer to the fly and human subsets of library DN as ‘putative *de novo* proteins’. In both cases, proteins were found to have weak, tissuespecific expression, and low-to-moderate signals of selection.

**Fig. 1.**
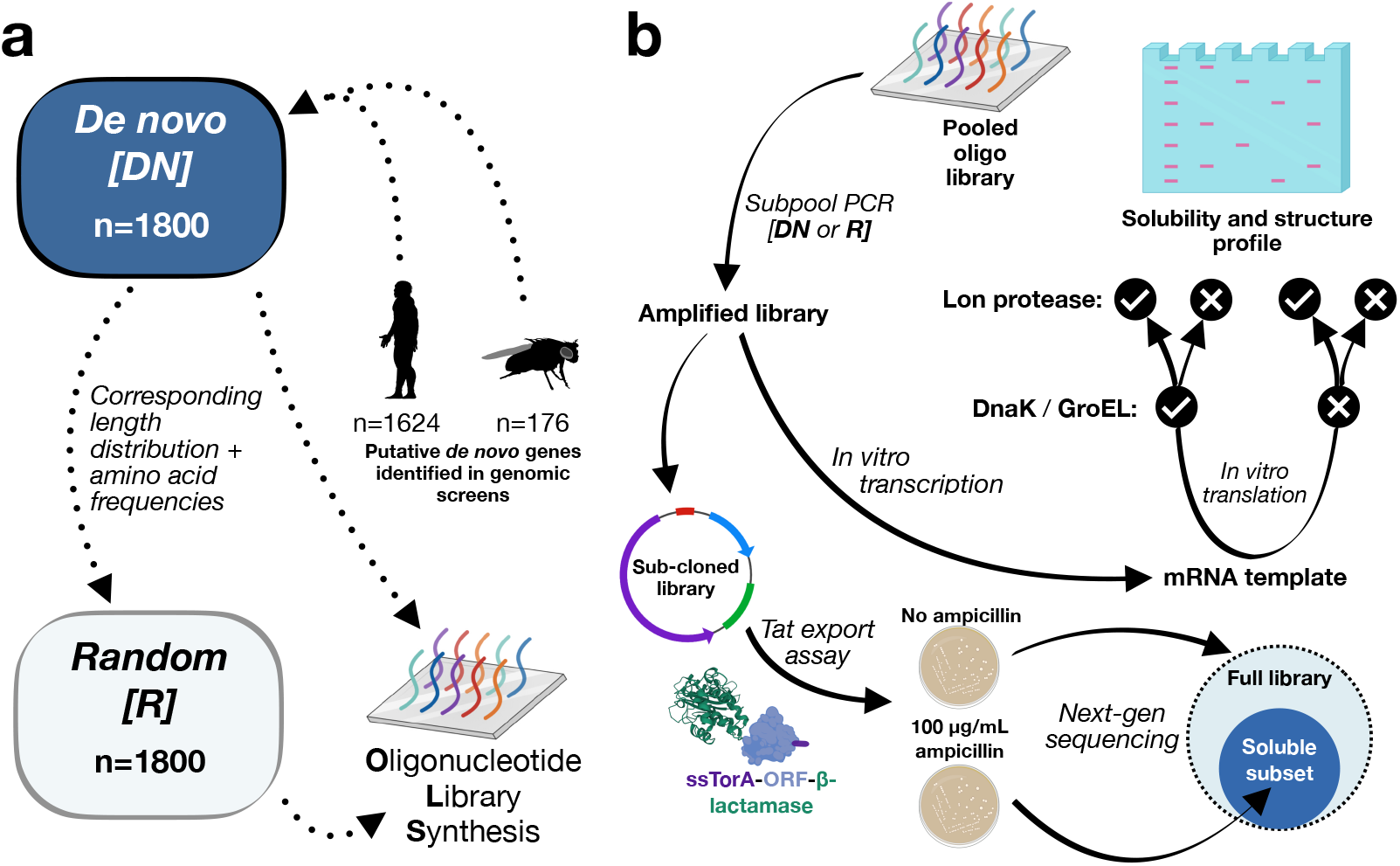
Library design, synthesis and experimental outline. **a)** Schematic illustrating the *in silico* design of libraries of *de novo* and un-evolved random-sequence proteins. A *de novo* library (DN) was built from putative *de novo* proteins identified in human and fly: subsequently, a library of unevolved random sequences (R) was designed to mirror the length and amino acid frequencies of library DN. The two libraries were synthesised by OLS ready for experimental study. **b)** Approaches used to profile solubility and structure content of each library. Following amplification, each library was expressed in a chaperone-assisted cell-free format, and a non-specific Lon protease was used to quantify structural content. In parallel, subcloned libraries were expressed in *E. coli* to screen for soluble and folded variants that did not disrupt periplasmic export.

Given the recent acquisition of these proteins and their apparent unevolved nature, it remains unclear how these novel proteins differ from ‘true’ random sequences if at all. For both human and fly sequences, various protein properties were predicted. Fly *de novo* proteins were compared to randomly sampled intergenic sequences without expression evidence, and found to have higher GC-content and ID. Human-specific ORFs identified by Dowling et al. (12), which make up the majority of the library DN, were not compared to a ‘more random’ set of sequences. However, they were found to have lower GC-content than conserved ORFs (i.e. ‘conservation level 5’, with exon overlap), but similar predicted ID. This discrepancy between GC-content and ID may be explained by the action of selection: either on newly emerged proteins, or over longer evolutionary timescales to shape the properties of highly-conserved ORFs.

To identify such selection towards a given biophysical property, a natural and feasible approach is to compare the set of putative *de novo* proteins to ‘true’ random controls and see if they differ. For this reason, a synthetic random library (R) was designed, with amino acid frequency and length distributions matched to library DN. Given that amino acid composition is a major determinant of all biophysical properties, the specification of library R should provide the most appropriate comparison; any differences in protein property between DN and R should be attributable to the specific residue ordering (and not compositional bias; see Fig. S2).

### *De novo* proteins are predicted to have highly similar properties to unnatural random sequences

Having designed libraries of putative *de novo* (library DN) and synthetic random proteins (library R) *in silico*, we next made a number of bioinformatic predictions of protein property. Figure 2 shows predictions for four relevant features. To put biophysical properties in context with those of conserved (i.e. ‘native-like’ proteins), predictions are compared to a length-matched subset of 3600 annotated human proteins. In all cases, predictions for DN and R are highly similar. Predictions of ID distribute similarly for all three classes (Fig. 2a), as does aggregation propensity (Fig. 2b). Comparison to annotated human proteins suggests reduced propensity for *Q*-helices in both libraries (Fig. 2c), but higher propensity for *β*-sheets (Fig. 2d). Accordingly, from primary sequence alone, libraries DN and R appear to have appropriate levels of hydrophobic and hydrophilic residues to form native-like structural content. For additional property predictions see Figure S1.

**Fig. 2.**
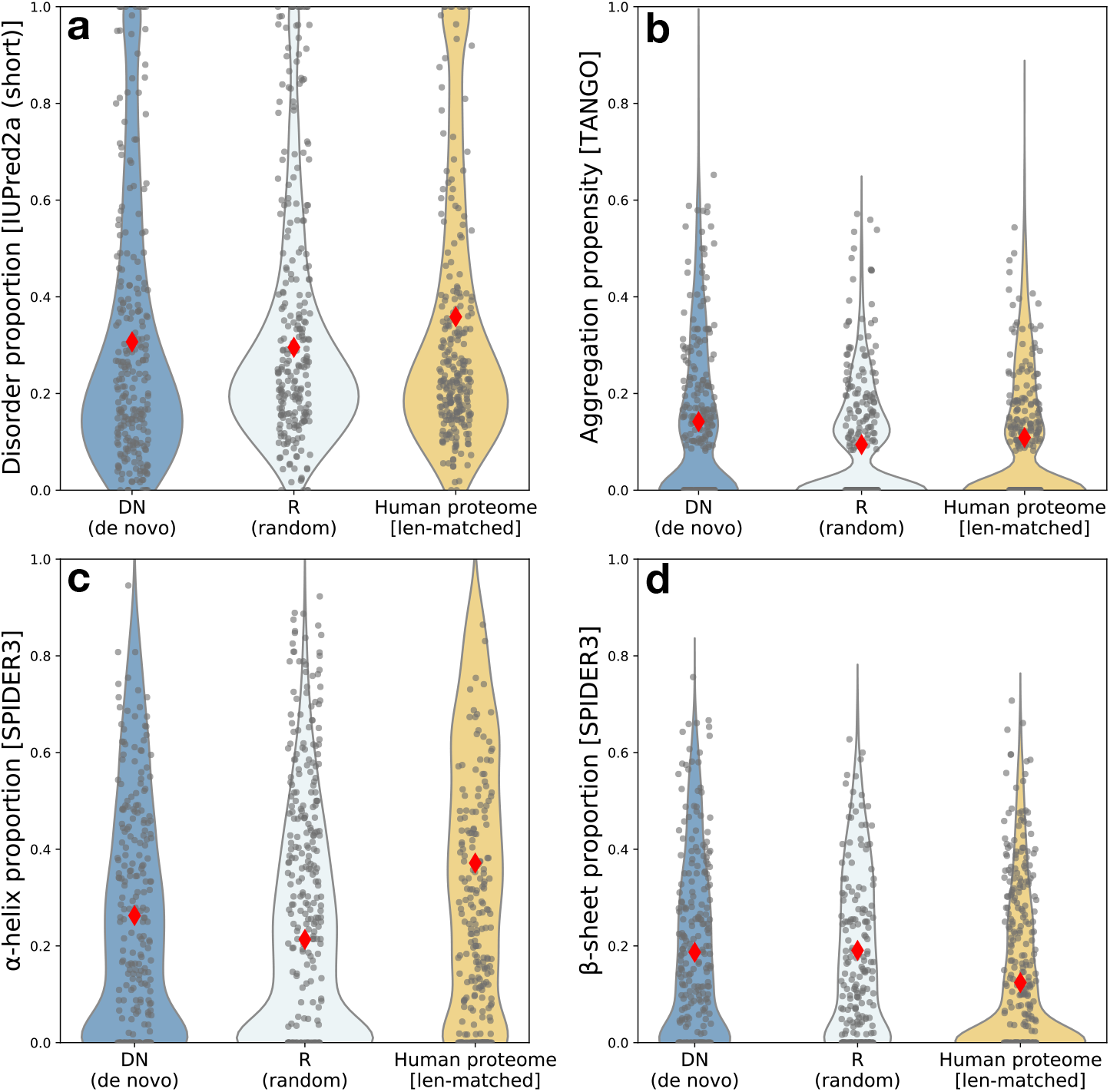
Biophysical predictions are similar for *de novo* and unevolved random sequences and suggest that both harbour high structural potential. Libraries DN (dark blue) and R (pale blue), designed to have matched length and amino acid frequencies, are predicted to have highly similar biophysical properties as expected. Comparison to a length-matched subset of the human proteome (yellow) shows broadly similar predictions, suggesting that native-like properties are present in, or at least evolutionarily accessible to, random sequence proteins. Diamonds indicate mean value of distributions, which are subsampled to 250 sequences for visualisation.

Prediction tools such as IUPred have been trained using the (relatively small) sets of proteins for which disorder or aggregation has been determined experimentally. Given the novel and unevolved nature of our libraries, we looked for a more generalisable predictor of structural content or stability. Learned embeddings have been described recently as a way to encode fundamental protein features learned over much larger regions of sequence space than have been experimentally characterised (33). For example, using UniRep embeddings as input, a linear model was shown to outperform Rosetta total energy predictions when trained on protease sensitivity data (34, 35).

Prior to an experimental protease assay (see following sections), we implemented this predictive model to generate protease stability scores for each library. As shown in Figure S3, we find libraries DN and R have highly similar predictions. The control set of annotated human proteins are predicted to be marginally more stable on average. However, scores broadly overlapped with those for the DN and R.

Derived from trypsin-based proteolysis data, the stability values predicted here are expected to correlate with total structure content and globularity. Accordingly, together with secondary structure predictions (Fig. 2), both libraries appear to have potential for structural content similar to that of conserved proteins. While *de novo* proteins may distribute to a particular region of protein sequence space – either due to selection or as a byproduct of their occurrence in a genome – library R is not similarly constrained. Instead, the similarity of all predictions for DN and R with those for conserved proteins appear to result from their similar amino acid compositions.

Aside from illustrating that all random sequences with appropriate amino acid composition may have structure forming potential, the predictions made here demonstrate that any difference between this set of putative *de novo* proteins and their unevolved random counterparts are indistinguishable computationally. This hypothesis is entirely plausible, but testing it computationally relies on the accuracy of the predictors used; predictors which may not be sensitive to small differences, especially when compositional biases are removed. For this reason, we next sought to validate these predictions experimentally.

### A cell-based protein export assay identifies soluble library members

Following *in silico* design, the libraries DN and R were synthesised as an oligonucleotide pool (Fig. 1a). *De novo* and random subpools were PCR amplified from this pool and used as a starting point for subsequent experimental work. We first used a twin-arginine export quality assay, which relies on translocation of *β*-lactamase via the twin-arginine translocation (Tat) pathway, to screen for soluble members of each library (36). This assay is implemented by sub-cloning each library to a vector encoding an N-terminal secretion-signal and a C-terminal *β*-lactamase (construct illustrated in Fig. 1b). Upon expression of the resulting fusion constructs in *E. coli,* successful export of the fused *β*-lactamase can be detected by colony formation on ampicillin plates. Ampicillin can therefore be used to select for library members that do not interfere with translocation. Twin-arginine export assay was previously shown to select for soluble target protein (37) and remove gene synthesis errors (38). We here use the assay to select for (and subsequently identify by sequencing) the soluble subsets of each library that do not result in aggregation of *β*-lactamase fusion proteins.

Selection of libraries DN and R on ampicillin, followed by NGS-based quantification of library diversity (i.e. the number of unique sequences represented), allows identification of soluble subsets of each library (and additionally an assessment of library quality: Figs. S4 & S5). Figure 3 shows the results selection on 100 *μ*g/mL ampicillin (the highest concentration assayed). When plated without ampicillin at 30 °C (Fig. 3a), 80-85% of theoretical library diversity was identified above a threshold of 100 reads-per-million (DN; 81%, 1452/1800. R; 83%, 1501/1800; for read-count distributions see Fig. S4). Post-selection on ampicillin, the fraction of the library identified by sequencing dropped to 43% and 34% for libraries DN and R, respectively. This indicates that the majority of both libraries are insoluble when expressed recombinantly in *E. coli*. Aggregation of *de novo* proteins expressed recombinantly has been noted previously and is consistent with this result (6).

**Fig. 3.**
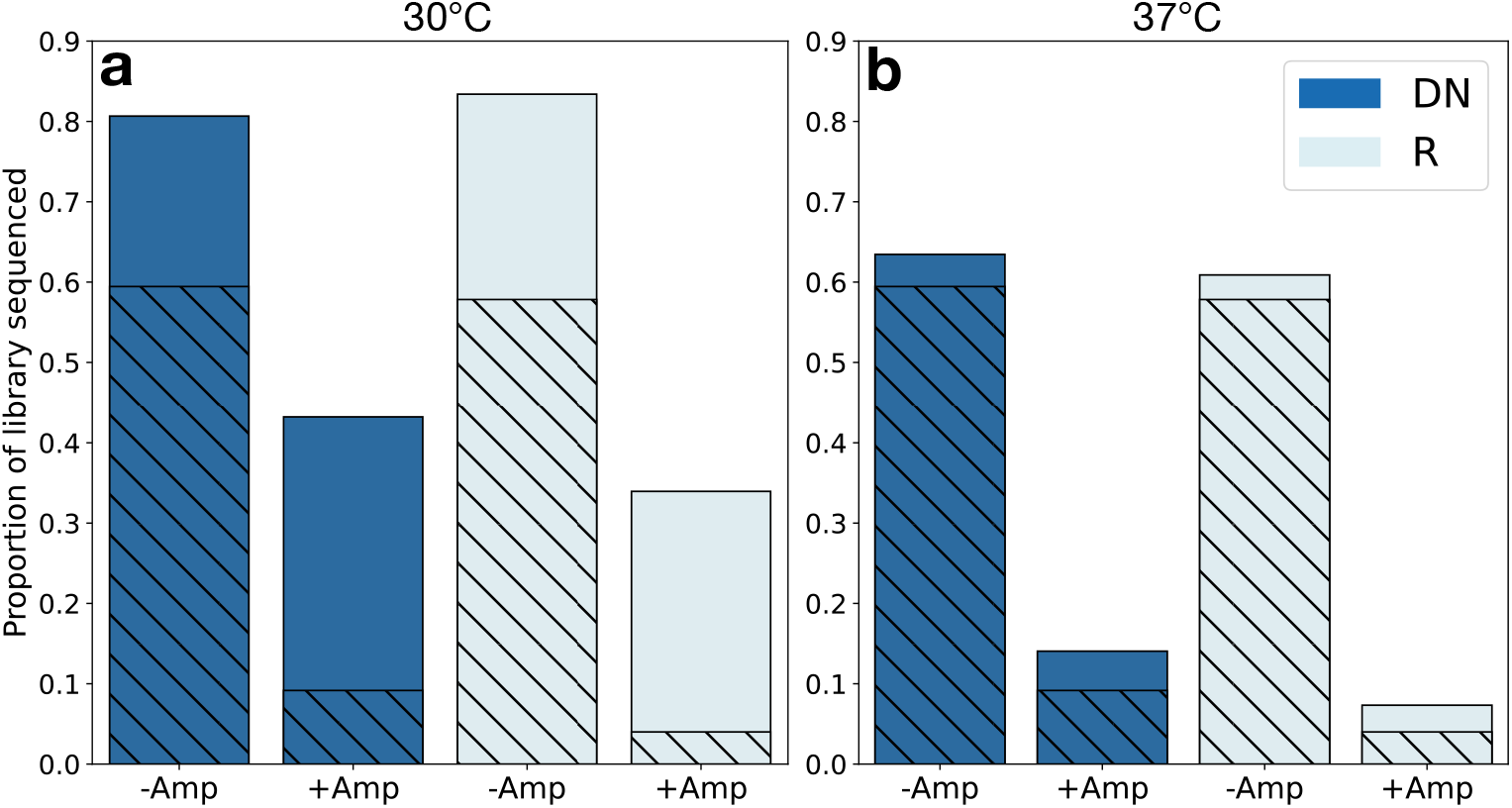
A cell-based assay identifies subsets of each library with potential for soluble expression. NGS of input (plated without ampicillin) and selected libraries (+100 *μ*g/mL ampicillin) allows quantification of changes in library diversity following twin-arginine export assay. **a)** Libraries DN and R had 43% and 34% representation following selection at 30 °C. **b)** At 37 °C, surviving fractions dropped to 14% (DN) and 7% (R), indicative of higher solubility for the putative *de novo* library. The majority of surviving variants at 37 °C also survived at 30 °C; hatched bars indicate shared representation at each temperature.

Repeating the same assay at 37 °C (Fig. 3b), we found overall lower diversity on pre-selection plates (DN; 63%, 1142/1800. R; 61%, 1050/1800). However, a greater relative drop in representation was seen upon ampicillin selection, to 14% and 7% for libraries DN and R, respectively. A greater efficacy of selection for solubility at 37 °C is consistent with greater overexpression at 30 °C – and could also indicate the presence of slow folders which are less able to avoid aggregation at increased temperatures. Interestingly, the trend for putative *de novo* proteins to have higher solubility on average than synthetic random sequences held at both temperatures, and was also consistent when split by human and fly subsets (see Fig. S6). Furthermore, many of the sequences selected at 37 °C were also selected at 30 °C and may represent the most ‘robustly soluble’ sequences.

### Putative *de novo* proteins may have higher intrinsic solubility than unevolved random controls

To further investigate the properties of our putative *de novo* and true random sets, libraries were expressed in a cell-free format using a reconstituted *E. coli* expression system including transcriptional and translational machinery. Cell-free (in vitro) recombinant expression has two key benefits in this case: first, it allows tight control of expression conditions and control of cofactor concentrations, and second, it separates intrinsic target-protein behaviour (e.g. aggregation propensity) from the complex cellular milieu (39). Libraries were expressed in vitro with a C-terminal FLAG^®^-tag and target protein detected by Western blot (Fig. 4a). In addition to total yield (T), the subset of soluble library protein is isolated and loaded in adjacent lanes (S). Quantification of the intensity of the soluble:total lanes therefore provides an estimate of the fraction of soluble expression in a given sample.

**Fig. 4.**
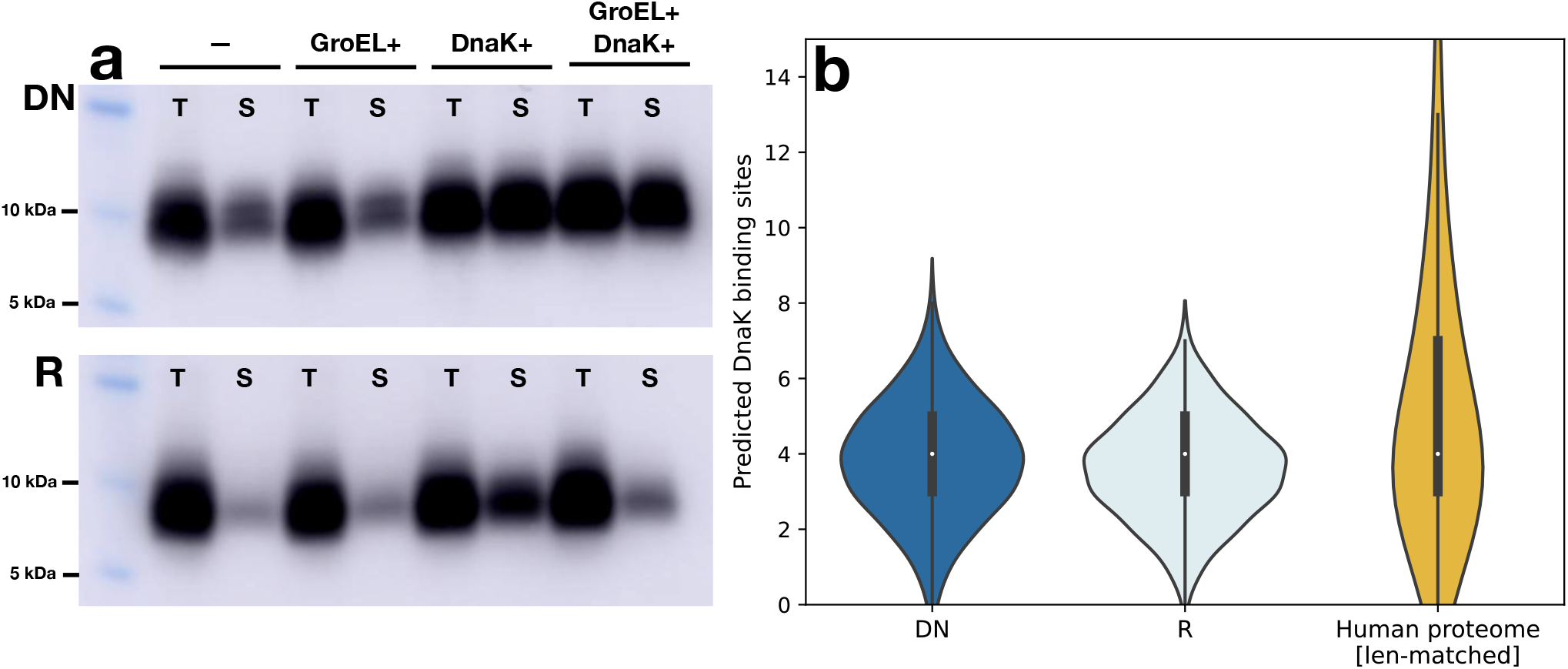
Cell-free expression indicates that a library of putative *de novo* proteins (DN) are more soluble than synthetic random sequences, and that DnaK solubilises both libraries equally well. **a)** Western blot showing total (T) and soluble (S) fractions of bulk library expression using reconstituted *E. coli* machinery in cell-free format (37°C). Library DN (top row) is marginally more soluble than library R (bottom row). Co-translational chaperone addition (DnaK, GroEL, or both) shows that GroEL has little effect but that DnaK solubilised both libraries. **b)** Bioinformatic prediction of the number of DnaK binding sites per sequence for libraries DN and R, with a length-matched set of annotated human proteins included for comparison.

Here, base expression (left panel) was compared to yield achieved in the presence of molecular chaperone systems added to the cell-free reaction (see Methods). GroEL/ES and DnaK systems were added co-translationally i.e, were present from the start of the reaction. As can be seen in Figure 4a, soluble protein makes up only a fraction of total expression in the absence of additional molecular chaperone. This was true for both the putative *de novo* proteins (top row) and the set of true random sequences (bottom row). Most notably, the soluble fraction inferred from blot intensity is consistent with the fraction of each library selected by the Twin-arginine assay at the same temperature (37°C). The same trend for DN to be slightly more soluble than R is also seen here.

Upon addition of GroEL/ES system (‘GroEL+’), no major difference in soluble yield was seen for either library. However, upon DnaK addition (‘DnaK+’) both libraries were highly solubilised (seen by intensity in lane S being close to that in lane T). When both DnaK and GroEL/ES systems were added, the improved solubility was maintained for library DN. However, for library R, addition of GroEL/ES appeared to counteract the effect of DnaK and solubility dropped closer to basal levels.

The difference seen between GroEL/ES and DnaK may be explained by their differing mechanisms; although GroEL has been shown to interact with random proteins (40), its more involved mechanism may require a greater degree of substrate adaptation. Figure 4b shows predicted DnaK binding sites for each library, compared to the subset of length-matched annotated human proteins. Library sequences are predicted to have on average four regions for which DnaK should have high affinity (short hydrophobic regions with positively charged residues) (Fig. 4b). This is comparable to the prediction for conserved proteins, which may help explain why DnaK is effective and acts similarly for libraries DN and R (~3-fold solubility increase).

### Proteolytic assay identifies large amount of undegradable protein for *de novo* and truly random sequences

Having seen consistent trends for the solubility of each library when expressed in *E. coli* as fusion constructs, and in a cell-free format, we next investigated the structural content using a Lon-based proteolytic assay (23, 41). Using the same cell-free expression system (see Fig. 4a), Lon protease was added to reaction mixtures. Lon’s preference for non-specific cleavage of exposed hydrophobic regions means that it causes the greatest amount of degradation for IDP-like proteins, and in general for proteins with lower structural propensity.

Figures 5a and 5b show representative blots for libraries DN and R, respectively, with addition of DnaK and Lon protease to cell-free reaction mixtures. Quantification of blot intensity over replicate blots allows an estimation of the degradable fractions of each library with respect to solubility (see Methods for more details). This is illustrated in Figure 5c, with soluble fractions (green hues) split by degradability (dark blue; soluble/undegraded, pale blue; soluble/degraded). The degraded and undegraded fractions of insoluble yield can also be inferred in this way (dark yellow; insoluble/undegraded, light yellow; insoluble/degraded).

**Fig. 5.**
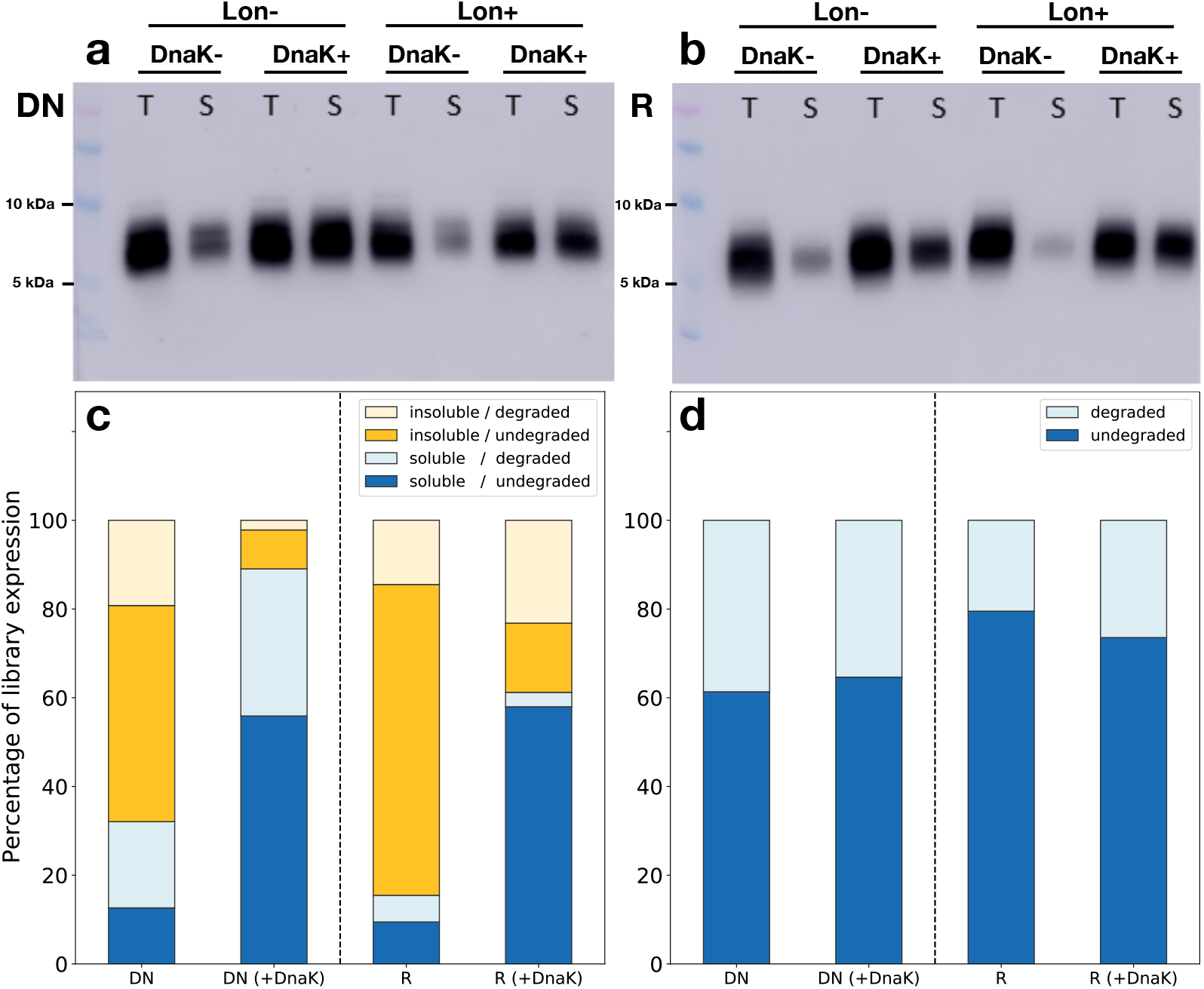
Quantification of degraded library fractions following cell-free expression in the presence of Lon protease. Total (T) and soluble (S) expression with co-translational (37°C) addition of DnaK and/or Lon protease: representative Western blots shown for libraries DN **(a)** and R **(b)**. Non-specific cleavage of hydrophobic regions by Lon protease results in preferential degradation of disordered proteins, with a visible net reduction in yield for Lon+ samples. **c)** Quantification of degraded fractions with respect to solubility reveals a greater IDP-like (soluble/degraded) fraction for putative *de novo* proteins vs. ‘true’ random sequences. DnaK addition, however, results in a greater increase in the soluble/undegraded fraction than the IDP-like fraction (for both DN and R). **d)** Summary of degraded vs undegraded fractions, regardless of solubility (sum of dark and light bars in (**c**), respectively). Library R is marginally less degradable than DN, suggesting slightly higher structural propensity.

As can be seen in Figs. 5a and 5b, addition of Lon protease causes a reduction in both the total yield and that of the soluble subset (where degradation is most visible). The fact that some soluble protein remains undegraded points to a degree of structural content even for the soluble fraction. In other words, a fraction of both the *de novo* and true random proteins have soluble expression, not all of which consists of IDP-like proteins (i.e. soluble and disordered). Quantifying this in Figure 5c shows that, considering only the soluble fraction, library DN has a greater proportion of these IDP-like proteins than library R, where less of the soluble fraction was degraded than not. In the insoluble fraction, for both libraries the majority of protein is inferred as undegradable. We suggest that this corresponds to insoluble proteins with above-average structural potential.

With addition of the DnaK, the same solubility increase as before (see Fig 4a) was seen. Comparing library DN to its no-DnaK reference suggests that, DnaK has acted to prevent much of the soluble/undegraded fraction from converting to the insoluble/undegraded fraction. Similarly, DnaK appears to have prevented much of library R’s soluble/undegradable fraction from aggregating. However, solubilisation of library R does not appear to result in a concurrent increase in the soluble/degraded fraction (IDP-like). This may be best explained by the overall lower degradation seen for library R. Combining soluble and insoluble fractions, library R can be seen to have higher apparent structural propensity compared to library DN (Fig. 5d).

## Discussion

Given an emerging picture of abundant structure and function within sequence space, an outstanding question is if *de novo* proteins differ from other classes of random protein. In other words: do *de novo* proteins occupy a privileged area of sequence space with respect to structure or function? Direct attempts to answer this question have so far not been made. Instead, experimental evidence from unnatural random sequence libraries have formed the basis for many hypotheses regarding *de novo* emergence. Further, direct investigation of *de novo* proteins has been limited to either computational prediction or experimental characterisation of individual proteins. Going beyond these studies, we assess a library of putative *de novo* proteins experimentally and compare their properties to a matched library of unevolved random sequences. In doing so, we show that recently emerged *de novo* proteins behave similarly to unevolved counterparts – but that the set of *de novo* proteins harbours a larger fraction of soluble and protease-sensitive sequences.

Recent improvements in DNA synthesis technology have made it feasible to generate large libraries of high-fidelity sequences. Using oligonucleotide library synthesis (OLS), it is possible to investigate proteins in high-throughput by direct specification of their coding sequences. We focus on short *de novo* proteins (<66 aa) that we previously identified in human and fly, which can be synthesised directly in a single oligonucleotide. However, multiplex gene synthesis also makes this approach applicable to longer proteins specified over multiple oligos (38, 42). Libraries generated in this way should ultimately allow coupling of computational identification and high-throughput investigation of diverse protein sequences.

Having designed a library of 1800 random sequences (R) to have matched amino-acid frequencies and lengths as a set of 1800 putative *de novo* sequences (DN), we ran primary-sequence based predictions for a number of biophysical properties. Given that all computational predictions are highly similar between the two libraries, a possible conclusion is that our library of *de novo* proteins is generally close to the set of synthetic random sequences, and that their shared biophysical propensities result from their matched amino acid compositions. However, the reliability of predictions for random-type proteins remains ambiguous, given that it is only possible to validate prediction tools on well characterised proteins which are typically well conserved. Furthermore, the predictors rely heavily on sliding-window assessments of sequence composition which could struggle to differentiate DN and R. In light of this, experimental characterisation remains critical to any conclusions regarding this class of proteins; a step that has until now not been attempted for more than a handful of *de novo* proteins.

We first assessed solubility of our libraries using a twin-arginine export quality assay (36), shown to select for soluble and folded proteins (38). Sequencing of libraries DN and R after selection showed that at least one third of each library (43% and 34%, respectively) has potential for soluble expression at 30 °C. Interestingly, computationally predicted properties did not correlate with those sequences most enriched by selection (i.e. the most soluble variants). Any distinguishing properties of these sequences were therefore not captured by computational tools, further highlighting the need for experimental characterisation.

Next, we expressed each library in cell-free format using reconstituted *E. coli* expression apparatus. Given that the putative *de novo* proteins were sourced from human and fly, cell-free expression allows separation of the inherent biophysical properties of each library and the unnatural *E. coli* cellular environment. In addition, the cell-free format enables systematic changes to expression conditions – including addition of molecular chaperones to aid solubility, or proteases to assess protein stability. In the absence of chaperones, we found putative *de novo* proteins to have higher solubility than their unevolved random counterparts (~30% soluble vs ~15%). This trend is in agreement with the the twin-arginine export assay, with a higher fraction of the *de novo* library having soluble potential. The higher solubility of putative *de novo* proteins may reflect their exposure to selection; avoidance of aggregation has been suggested as a key selective pressure on novel proteins (15). Despite their recent emergence, and typically low and tissue-specific expression, selection may have shaped the properties of these sequences to some degree.

We next tested the effect of two chaperone systems, GroEL and DnaK, on the expression of each library. While GroEL had no effect on solubility or overall expression, DnaK increased the soluble fraction of both libraries by ~’-3-fold. This resulted in soluble fractions of ~90%. (DN) and ~60% (R), most likely due to DnaK having similar effectiveness on both libraries and preventing approximately equal amounts of protein from forming insoluble aggregates. The effectiveness of DnaK on random proteins was demonstrated recently (23). Confirming this result for putative *de novo* proteins indicates that DnaK (or its eukaryotic homolog Hsp70) may be essential for avoidance of aggregation in the early stages of protein evolution.

Finally, to probe the structural content of each library, we included Lon protease in the cell-free expression system (41). By preferentially cleaving exposed hydrophobic regions of unstructured proteins, Lon degradation correlates with intrinsic disorder (43). A Lon-based method was recently used to probe random-sequence libraries of different amino acid compositions (23), identifying a significant proportion of the soluble fraction of each library to be resistant to degradation. In addition, increasing solubility with DnaK also had a small effect on the fraction of non-degradable protein. While the precise fractions of degraded protein for each condition should be interpreted with care, in both cases over 50% of soluble protein was not degraded by Lon upon DnaK addition. A subset of each library may therefore harbour structural elements that interfere with cleavage, in agreement with findings that structure is abundant in sequence space (23). However, the low resolution of the Lon-assay prevents differentiation of different forms of structural elements, such as oligomeric or molten globule. Interestingly, we find 10-20% higher degradation for putative *de novo* proteins compared to synthetic random sequences, in agreement with our earlier report showing that unevolved sequences with less structural content are more soluble upon expression in *E. coli* (22).

Although putative *de novo* proteins appear marginally more soluble than synthetic random proteins, both show sensitivity to molecular chaperones. Similarly, while a subset of both libraries may harbour structural content, putative *de novo* proteins appear to contain more disordered regions, in correlation with their higher solubility. We note that our study is limited to short proteins of a specific composition and GC-content distribution. While the results presented here transcend earlier computational analyses and studies of single *de novo* proteins, we note that it is not possible to ultimately prove any instance of *de novo* emergence and there remains a degree of uncertainty about the true origin of the putative *de novo* proteins studied here. Some of the putative *de novo* set, in particular those from *H. sapiens,* may in fact be transient short-lived proto-genes which have not yet assumed critical cellular roles (but are nonetheless evolutionarily highly relevant; see Keeling et al. (44)).

In summary, we suggest that *de novo* proteins are not especially privileged among random sequences, and that the propensity for structure across sequence space may be key to the feasibility of *de novo* emergence. However, our findings of higher solubility for putative *de novo* proteins are consistent with early selection pressure to avoid aggregation. To corroborate this finding, larger numbers of *de novo* proteins drawn from diverse genomic backgrounds should be characterised in future efforts.

## DATA AVAILABILITY

Library sequences and code used for library design can be found at: https://zivgitlab.uni-muenster.de/agebb/de-novo/de_novo_lib

## ACKNOWLEDGEMENTS

This project has received funding from the European Union’s Horizon 2020 research and innovation programme under the Marie Skłodowska-Curie grant agreement No 722610. KH, AL and MA were funded by Volkswagen Stiftung (VWF), grant code 98183 to KH and EBB. KH, FB and VT were additionally funded by Primus grant PRIMUS/20/SCI/012 from Charles University. MA received funding from a DAAD Research Scholarship for doctoral students. Open Access funding provided by Projekt DEAL. Figure 1 was created with BioRender.com.

## Methods

### Library sequence selection

To study the properties of *de novo* and random-sequence proteins experimentally, two libraries were first designed *in silico*. In prior work, we identified large sets of putative *de novo* proteins which appear to have emerged from previously non-coding DNA. To build a *de novo* library (DN), 1800 proteins were selected from two studies identifying *de novo* genes in fly (n=176) (13), and newly-transcribed human ORFs (n=1624) (12) (‘conservation level 0’ in Dowling et al. (2020), excluding ORFs with exon-overlap). A library of 1800 unevolved random-sequence proteins (R) was then generated synthetically by sampling amino acids using the frequency distribution of library DN. Sequence lengths were also matched to those of library DN, so that library R had identical length and amino acid composition to library DN.

### Oligonucleotide pool design

Libraries DN and R were synthesised as a SurePrint oligonucleotide pool by Agilent (DE). Oligonucleotides were specified to include NdeI and XhoI restriction sites 5’- and 3’ to the CDS for downstream cloning. Additionally, 15 bp primer sites were added up- and downstream of the restriction sites to allow libraries DN and R to be PCR amplified separately from the oligo pool. The DnaChisel package (45) was used to codon optimise CDSs for protein expression in *E. coli*, while avoiding introduction of undesired restriction sites and homopolymer repeats of 5 bp or longer. Starting from desired amino acid sequences, we selected the highest frequency codon according to *E. coli* K12 frequencies (http://www.kazusa.or.jp/codon), and DnaChisel’s ‘harmonized Relative Codon Adaptiveness’ implementation was used to replace rare codons (46). Code to generate optimised oligo pools was used here as follows to select and optimise the 1800 longest compatible open reading frames (ORFs) from a list of human and fly *de novo* ORFs:

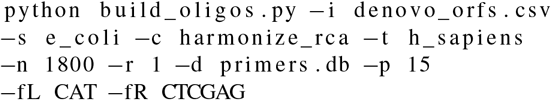

### Prediction of protein properties

Intrinsic structural disorder and globularity were calculated using IUPred2a (47); secondary structure, Phi and accessible surface area (ASA) were predicted using SPIDER3 (48); aggregation propensity was predicted using TANGO (49); isoelectric point (IEP) was predicted using EMBOSS pepstats (50); and grand average of hydropathy (GRAVY) index was calculated using CodonW (51). To predict stability scores, we used an implementation of UniRep (35, 52) to generate sequence embeddings of size 1900, and trained a sparse linear model (Lasso least-angle regression with 10-fold cross-validation) on a dataset of *de novo*-designed proteins with experimentally determined stability scores (34), as described by Alley et al. (35). As a comparison for predictions, 3600 annotated human proteins (Ensembl 97 *H. sapiens* proteome) were selected by random sampling of an equal-length protein for each member of library DN. DnaK binding sites were predicted using the ChaperISM suite (v1) in quantitative mode with default settings (53).

### Twin-arginine export quality assay

To screen for soluble proteins, libraries were expressed as fusions with an N-terminal Tat secretion signal (ssTorA) and a C-terminal *β*-lactamase. Misfolding or aggregation of the target ORF should prevent secretion of the construct to the *E. coli* periplasm, allowing selection by plating on increasing concentrations of ampicillin. Libraries DN and R were PCR-amplified separately from the oligonucleotide pool, with primers introducing EcoRI and BamHI restriction sites. After restriction cloning to pSALECT-EcoBam (Addgene plasmid 59705), libraries were transformed by electroporation to E. cloni 10G SOLO cells (Lucigen). Whole transformations were plated on LB-agar + 25 *μ*g/mL chloramphenicol and grown overnight. Libraries were then scraped from plate into LB medium to make glycerol stocks adjusted to have the same OD600. Stocks were kept at −80 °C and used for all subsequent plating assays. The assay involved plating equal volumes of glycerol stock on LB-agar supplemented with either: 25 *μ*g/mL chloramphenicol, 25 *μ*g/mL chloramphenicol and 4 *μ*g/mL ampicillin, or 25 *μ*g/mL chloramphenicol and 100 *μ*g/mL ampicillin. After incubation overnight at 30 °C, plates were scraped into PBS and plasmid isolated (GeneJET Plasmid Miniprep Kit, Thermo Scientific). Primers encoding 8-bp 5’ and 3’ barcodes were used to amplify samples from each condition.

### Next-generation sequencing

Amplicons from Twin-arginine export assay conditions were purified, combined in equimolar amounts, and amplicon size distribution (270-350 bp) verified by capillary electrophoresis. Amplicons were subsequently sequenced using an Illumina MiSeq platform. Reads were merged, trimmed and filtered to remove low quality reads using the fastp suite (54). The cutadapt suite (55) was used for read demultiplexing, and reads were then mapped to CDS sequences of libraries DN and R using the Burrows-Wheeler Alignment (BWA) MEM algorithm (56). SAMtools was used for conversion to SAM file format, sorting and indexing (57). Finally, reads mapped to each variant were counted using HTSeq (58). Read counts were converted to reads-per-million-reads (RPM) values (per plating condition) to control for sequencing depth, and sequences were subsequently filtered using a threshold of 100 RPM to remove those with very low abundance (i.e. <0.01% of reads in a given sample).

### Cell-free expression and Lon proteolytic assay

Both protein libraries were produced in a cell-free expression system to evaluate their solubility, response to chaperones and structural content (using proteolysis resistance) in a cell-like environment. Expression from mRNA templates was carried out in *E. coli* reconstituted cell-free system and solubility was assessed by centrifugation to separate soluble fraction, followed by quantitative Western blot. Bacterial Lon protease preferentially cleaves unstructured proteins and was added to the reactions to investigate proteolytic resistance potential of the protein libraries (23).

First, library subpools were PCR-amplified to introduce EcoRI and BamHI restriction sites, subcloned into pET24a+ vector modified to encode a C-terminal FLAG^®^-tag and electroporated into *E. cloni* 10G cells (Lucigen, USA). Cells were grown overnight at 37 °C on LB-agar + 50 *μ*g/mL kanamycin plates and transformants scraped for plasmid DNA isolation. The region containing the T7 promoter, library sequence and terminator was PCR amplified to serve as template for *in vitro* transcription (NEB HiScribe T7 kit, USA). The PUREfrex 2.0 system (GeneFrontier Corporation, Japan) was used for *in vitro* translation. The reactions were mixed as per protocol to final volume 10 *μ*L with addition of 0.05% Triton X-100 and incubated at 37 °C for 2 h. To assess the effect of molecular chaperones on the soluble yield of protein expression, reactions were supplemented with DnaK or GroE mix (GeneFrontier Corporation, Japan), to final concentration of 5 *μ*M DnaK, 1 *μ*M DnaJ and GrpE, 0.1 *μ*M GroEL and 0.2 *μ*M GroES. For proteolytic resistance assay, purified Lon protease was added co-translationally at 0.1 *μ*M working concentration.

Following production all reactions were halted by adding 40 *μ*L of puromycin buffer (300 *μ*M puromycin, 50 mM Tris, 100 mM NaCl, 100 mM KCl, pH 7.5) and incubating at 30 °C for 30 min. Next, 5 *μ*L of such mixture was processed for SDS-PAGE serving as the total (T) fraction of expression, while the rest was centrifuged (21,000 x g, 30 min, 21 °C). Soluble (S) fraction was collected by taking 5 *μ*L of the supernatant. Finally, three technical replicates for each sample were analyzed by SDS-PAGE and Western blot using Anti-FLAG^®^ antibody (Sigma-Aldrich Monoclonal ANTIFLAG^®^ M2-Peroxidase (HRP) antibody, A8592). Images were quantified using ImageJ (U. S. National Institutes of Health, USA).

## Bibliography

1. Jonathan F Schmitz and Erich Bornberg-Bauer. Fact or fiction: updates on how proteincoding genes might emerge de novo from previously non-coding DNA. F1000Research, 6: 57, January 2017. ISSN 2046-1402. doi: 10.12688/f1000research.10079.1.

2. Nikolaos Vakirlis, Omer Acar, Brian Hsu, Nelson Castilho Coelho, S. Branden Van Oss, Aaron Wacholder, Kate Medetgul-Ernar, Ray W. Bowman, Cameron P. Hines, John Iannotta, Saurin Bipin Parikh, Aoife McLysaght, Carlos J. Camacho, Allyson F. O’Donnell, Trey Ideker, and Anne-Ruxandra Carvunis. De novo emergence of adaptive membrane proteins from thymine-rich genomic sequences. Nature Communications, 11(1):781, February 2020. ISSN 2041-1723. doi: 10.1038/s41467-020-14500-z. Number: 1 Publisher: Nature Publishing Group.

3. Stephen Branden Van Van Oss and Anne-Ruxandra Carvunis. De novo gene birth. PLOS Genetics, 15(5):e1008160, May 2019. ISSN 1553-7404. doi: 10.1371/journal.pgen.1008160.

4. Erich Bornberg-Bauer, Klara Hlouchova, and Andreas Lange. Structure and function of naturally evolved de novo proteins. Current Opinion in Structural Biology, 68:175–183, June 2021. ISSN 0959-440X. doi: 10.1016/j.sbi.2020.11.010.

5. Anna M. Gubala, Jonathan F. Schmitz, Michael J. Kearns, Tery T. Vinh, Erich Bornberg-Bauer, Mariana F. Wolfner, and Geoffrey D. Findlay. The goddard and saturn genes are essential for Drosophila male fertility and may have arisen de novo. Molecular Biology and Evolution, January 2017. ISSN 1537-1719. doi: 10.1093/molbev/msx057.

6. Andreas Lange, Prajal H. Patel, Brennen Heames, Adam M. Damry, Thorsten Saenger, Colin J. Jackson, Geoffrey D. Findlay, and Erich Bornberg-Bauer. Structural and functional characterization of a putative de novo gene in Drosophila. Nature Communications, 12 (1):1667, March 2021. ISSN 2041-1723. doi: 10.1038/s41467-021-21667-6. Number: 1 Publisher: Nature Publishing Group.

7. Emily L. Rivard, Andrew G. Ludwig, Prajal H. Patel, Anna Grandchamp, Sarah E. Arnold, Alina Berger, Emilie M. Scott, Brendan J. Kelly, Grace C. Mascha, Erich Bornberg-Bauer, and Geoffrey D. Findlay. A putative de novo evolved gene required for spermatid chromatin condensation in Drosophila melanogaster. PLOS Genetics, 17(9):e1009787, September 2021. ISSN 1553-7404. doi: 10.1371/journal.pgen.1009787. Publisher: Public Library of Science.

8. Chen Xie, Cemalettin Bekpen, Sven Künzel, Maryam Keshavarz, Rebecca Krebs-Wheaton, Neva Skrabar, Kristian Karsten Ullrich, and Diethard Tautz. A de novo evolved gene in the house mouse regulates female pregnancy cycles. eLife, 8:e44392, August 2019. ISSN 2050-084X. doi: 10.7554/eLife.44392.

9. Dixie Bungard, Jacob S. Copple, Jing Yan, Jimmy J. Chhun, Vlad K. Kumirov, Scott G. Foy, Joanna Masel, Vicki H. Wysocki, and Matthew H. J. Cordes. Foldability of a Natural De Novo Evolved Protein. Structure, 25(11):1687–1696.e4, November 2017. ISSN 0969-2126. doi: 10.1016/j.str.2017.09.006.

10. Helle Tessand Baalsrud, Ole Kristian Tørresen, Monica Hongrø Solbakken, Walter Salzburger, Reinhold Hanel, Kjetill S. Jakobsen, and Sissel Jentoft. De Novo Gene Evolution of Antifreeze Glycoproteins in Codfishes Revealed by Whole Genome Sequence Data. Molecular Biology and Evolution, 35(3):593–606, March 2018. ISSN 0737-4038. doi: 10.1093/molbev/msx311. Publisher: Oxford Academic.

11. Claudio Casola. From De Novo to “De Nono”: The Majority of Novel Protein-Coding Genes Identified with Phylostratigraphy Are Old Genes or Recent Duplicates. Genome Biology and Evolution, 10(11):2906–2918, November 2018. doi: 10.1093/gbe/evy231.

12. Daniel Dowling, Jonathan F Schmitz, and Erich Bornberg-Bauer. Stochastic Gain and Loss of Novel Transcribed Open Reading Frames in the Human Lineage. Genome Biology and Evolution, 12(11):2183–2195, November 2020. ISSN 1759-6653. doi: 10.1093/gbe/evaa194.

13. Brennen Heames, Jonathan Schmitz, and Erich Bornberg-Bauer. A Continuum of Evolving De Novo Genes Drives Protein-Coding Novelty in Drosophila. Journal of Molecular Evolution, April 2020. ISSN 1432-1432. doi: 10.1007/s00239-020-09939-z.

14. Jonathan F. Schmitz, Kristian K. Ullrich, and Erich Bornberg-Bauer. Incipient de novo genes can evolve from frozen accidents that escaped rapid transcript turnover. Nature Ecology & Evolution, 2(10):1626–1632, October 2018. ISSN 2397-334X. doi: 10.1038/s41559-018-0639-7.

15. Annamária F. Ángyán, András Perczel, and Zoltán Gáspári. Estimating intrinsic structural preferences of de novo emerging random-sequence proteins: Is aggregation the main bottleneck? FEBS Letters, 586(16):2468–2472, 2012. ISSN 1873-3468. doi: https://doi.org/10.1016/j.febslet.2012.06.007. _eprint: https://febs.onlinelibrary.wiley.com/doi/pdf/10.1016/j.febslet.2012.06.007.

16. Nicolas Galtier, Camille Roux, Marjolaine Rousselle, Jonathan Romiguier, Emeric Figuet, Sylvain Glémin, Nicolas Bierne, and Laurent Duret. Codon Usage Bias in Animals: Disentangling the Effects of Natural Selection, Effective Population Size, and GC-Biased Gene Conversion. Molecular Biology and Evolution, 35(5):1092–1103, May 2018. ISSN 1537-1719. doi: 10.1093/molbev/msy015.

17. Walter Basile, Marco Salvatore, and Arne Elofsson. The classification of orphans is improved by combining searches in both proteomes and genomes. bioRxiv, page 185983, May 2019. doi: 10.1101/185983.

18. Jiří Vymětal, Jiří Vondrášek, and Klára Hlouchová. Sequence Versus Composition: What Prescribes IDP Biophysical Properties? Entropy, 21(7):654, July 2019. doi: 10.3390/e21070654.

19. Cristiano Chiarabelli, Jan W. Vrijbloed, Richard M. Thomas, and PierLuigi Luisi. Investigation of de novo Totally Random Biosequences, Part I. Chemistry & Biodiversity, 3(8):827–839, August 2006. ISSN 1612-1880. doi: 10.1002/cbdv.200690087.

20. Thomas H. LaBean, Tauseef R. Butt, Stuart A. Kauffman, and Erik A. Schultes. Protein Folding Absent Selection. Genes, 2(3):608–626, August 2011. ISSN 2073-4425. doi: 10.3390/genes2030608.

21. Jia-Feng Yu, Zanxia Cao, Yuedong Yang, Chun-Ling Wang, Zhen-Dong Su, Ya-Wei Zhao, Ji-Hua Wang, and Yaoqi Zhou. Natural protein sequences are more intrinsically disordered than random sequences. Cellular and molecular life sciences: CMLS, 73(15):2949–2957, August 2016. ISSN 1420-9071. doi: 10.1007/s00018-016-2138-9.

22. Vyacheslav Tretyachenko, Jiří Vymětal, Lucie Bednárová, Vladimír Kopecký, Kateřina Hofbauerová, Helena Jindrová, Martin Hubálek, Radko Souček, Jan Konvalinka, Jiří Vondrášek, and Klára Hlouchová. Random protein sequences can form defined secondary structures and are well-tolerated in vivo. Scientific Reports, 7(1):15449, November 2017. ISSN 2045-2322. doi: 10.1038/s41598-017-15635-8.

23. Vyacheslav Tretyachenko, Jiří Vymětal, Tereza Neuwirthová, Jiří Vondrášek, Kosuke Fu-jishima, and Klára Hlouchová. Structured proteins are abundant in unevolved sequence space. preprint, Synthetic Biology, August 2021.

24. Yuuki Hayashi, Hiroshi Sakata, Yoshihide Makino, Itaru Urabe, and Tetsuya Yomo. Can an Arbitrary Sequence Evolve Towards Acquiring a Biological Function? Journal of Molecular Evolution, 56(2):162–168, February 2003. ISSN 0022-2844, 1432-1432. doi: 10.1007/s00239-002-2389-y.

25. Anthony D. Keefe and Jack W. Szostak. Functional proteins from a random-sequence library. Nature, 410(6829):715–718, April 2001. ISSN 0028-0836. doi: 10.1038/35070613.

26. C. A. Kaiser, D. Preuss, P. Grisafi, and D. Botstein. Many random sequences functionally replace the secretion signal sequence of yeast invertase. Science, 235(4786):312–317, January 1987. ISSN 0036-8075, 1095-9203. doi: 10.1126/science.3541205.

27. Michael Knopp, Jonina S. Gudmundsdottir, Tobias Nilsson, Finja König, Omar Warsi, Fredrika Rajer, Pia Ädelroth, and Dan I. Andersson. De Novo Emergence of Peptides That Confer Antibiotic Resistance. mBio, 10(3):e00837–19, June 2019. ISSN 2150-7511. doi: 10.1128/mBio.00837-19.

28. Michael Knopp, Arianne M. Babina, Jónína S. Gudmundsdóttir, Martin V Douglass, M. Stephen Trent, and Dan I. Andersson. A novel type of colistin resistance genes selected from random sequence space. PLOS Genetics, 17(1):e1009227, January 2021. ISSN 1553-7404. doi: 10.1371/journal.pgen.1009227. Publisher: Public Library of Science.

29. Rafik Neme, Cristina Amador, Burcin Yildirim, Ellen McConnell, and Diethard Tautz. Random sequences are an abundant source of bioactive RNAs or peptides. Nature Ecology &Evolution, 1(6):0127, April 2017. ISSN 2397-334X. doi: 10.1038/s41559-017-0127.

30. D. D. Axe, N. W. Foster, and A. R. Fersht. Active barnase variants with completely random hydrophobic cores. Proceedings of the National Academy of Sciences of the United States of America, 93(11):5590–5594, May 1996. ISSN 0027-8424.

31. Fa-An Chao, Aleardo Morelli, John C. Haugner Iii, Lewis Churchfield, Leonardo N. Hag-mann, Lei Shi, Larry R. Masterson, Ritimukta Sarangi, Gianluigi Veglia, and Burckhard Seelig. Structure and dynamics of a primordial catalytic fold generated by *in vitro* evolution. Nature Chemical Biology, 9(2):81–83, February 2013. ISSN 1552-4469. doi: 10.1038/nchembio.1138.

32. Asao Yamauchi, Toshihiro Nakashima, Nobuhiko Tokuriki, Masato Hosokawa, Hideki Nogami, Shingo Arioka, Itaru Urabe, and Tetsuya Yomo. Evolvability of random polypeptides through functional selection within a small library. Protein Engineering, 15(7):619–626, July 2002. ISSN 0269-2139.

33. Kevin K Yang, Zachary Wu, Claire N Bedbrook, and Frances H Arnold. Learned protein embeddings for machine learning. Bioinformatics, 34(15):2642–2648, August 2018. ISSN 1367-4803. doi: 10.1093/bioinformatics/bty178.

34. Gabriel J. Rocklin, Tamuka M. Chidyausiku, Inna Goreshnik, Alex Ford, Scott Houliston, Alexander Lemak, Lauren Carter, Rashmi Ravichandran, Vikram K. Mulligan, Aaron Chevalier, Cheryl H. Arrowsmith, and David Baker. Global analysis of protein folding using massively parallel design, synthesis, and testing. Science, 357(6347):168–175, July 2017. ISSN 0036-8075, 1095-9203. doi: 10.1126/science.aan0693.

35. Ethan C. Alley, Grigory Khimulya, Surojit Biswas, Mohammed AlQuraishi, and George M. Church. Unified rational protein engineering with sequence-based deep representation learning. Nature Methods, 16(12):1315–1322, December 2019. ISSN 1548-7105. doi: 10.1038/s41592-019-0598-1. Number: 12 Publisher: Nature Publishing Group.

36. Adam C. Fisher, Woojin Kim, and Matthew P. Delisa. Genetic selection for protein solubility enabled by the folding quality control feature of the twin-arginine translocation pathway. Protein Science, 15(3):449–458, 2006. ISSN 1469-896X. doi: https://doi.org/10.1110/ps.051902606. _eprint: https://onlinelibrary.wiley.com/doi/pdf/10.1110/ps.051902606.

37. Hyung-Kwon Lim, Thomas J. Mansell, Stephen W. Linderman, Adam C. Fisher, Michael R. Dyson, and Matthew P. DeLisa. Mining mammalian genomes for folding competent proteins using Tat-dependent genetic selection in Escherichia coli. Protein Science, 18(12):2537–2549, 2009. ISSN 1469-896X. doi: 10.1002/pro.262. _eprint: https://onlinelibrary.wiley.com/doi/pdf/10.1002/pro.262.

38. Timothy H.-C. Hsiau, David Sukovich, Phillip Elms, Robin N. Prince, Tobias Stritmatter, Paul Ruan, Bo Curry, Paige Anderson, Jeff Sampson, and J. Christopher Anderson. A Method for Multiplex Gene Synthesis Employing Error Correction Based on Expression. PLOS ONE, 10 (3):e0119927, March 2015. ISSN 1932-6203. doi: 10.1371/journal.pone.0119927. Publisher: Public Library of Science.

39. Tatsuya Niwa, Bei-Wen Ying, Katsuyo Saito, WenZhen Jin, Shoji Takada, Takuya Ueda, and Hideki Taguchi. Bimodal protein solubility distribution revealed by an aggregation analysis of the entire ensemble of Escherichia coli proteins. Proceedings of the National Academy of Sciences, 106(11):4201–4206, March 2009. ISSN 0027-8424, 1091-6490. doi: 10.1073/pnas.0811922106.

40. K. Aoki, F. Motojima, H. Taguchi, T. Yomo, and M. Yoshida. GroEL binds artificial proteins with random sequences. The Journal of Biological Chemistry, 275(18):13755–13758, May 2000. ISSN 0021-9258. doi: 10.1074/jbc.275.18.13755.

41. Tatsuya Niwa, Eri Uemura, Yuki Matsuno, and Hideki Taguchi. Translation-coupled protein folding assay using a protease to monitor the folding status. Protein Science, 28(7):1252–1261, 2019. ISSN 1469-896X. doi: 10.1002/pro.3624. _eprint: https://onlinelibrary.wiley.com/doi/pdf/10.1002/pro.3624.

42. Jason C. Klein, Marc J. Lajoie, Jerrod J. Schwartz, Eva-Maria Strauch, Jorgen Nelson, David Baker, and Jay Shendure. Multiplex pairwise assembly of array-derived DNA oligonucleotides. Nucleic Acids Research, 44(5):e43, March 2016. ISSN 0305-1048. doi: 10.1093/nar/gkv1177.

43. Laurence Van Melderen and Abram Aertsen. Regulation and quality control by Lon-dependent proteolysis. Research in Microbiology, 160(9):645–651, November 2009. ISSN 1769-7123. doi: 10.1016/j.resmic.2009.08.021.

44. Diane Marie Keeling, Patricia Garza, Charisse Michelle Nartey, and Anne-Ruxandra Carvunis. The meanings of ‘function’ in biology and the problematic case of de novo gene emergence. eLife, 8:e47014, November 2019. ISSN 2050-084X. doi: 10.7554/eLife.47014.

45. Valentin Zulkower and Susan Rosser. DNA Chisel, a versatile sequence optimizer. Bioinformatics, September 2020. doi: 10.1093/bioinformatics/btaa558.

46. Nico J. Claassens, Melvin F. Siliakus, Sebastiaan K. Spaans, Sjoerd C. A. Creutzburg, Bart Nijsse, Peter J. Schaap, Tessa E. F. Quax, and John van der Oost. Improving heterologous membrane protein production in Escherichia coli by combining transcriptional tuning and codon usage algorithms. PLOS ONE, 12(9):e0184355, September 2017. ISSN 1932-6203. doi: 10.1371/journal.pone.0184355. Publisher: Public Library of Science.

47. Bálint Mészáros, Gábor Erdos, and Zsuzsanna Dosztányi. IUPred2A: context-dependent prediction of protein disorder as a function of redox state and protein binding. Nucleic Acids Research, 46(Web Server issue):W329–W337, July 2018. ISSN 0305-1048. doi: 10.1093/nar/gky384.

48. Rhys Heffernan, Yuedong Yang, Kuldip Paliwal, Yaoqi Zhou, and Alfonso Valencia. Capturing non-local interactions by long short-term memory bidirectional recurrent neural networks for improving prediction of protein secondary structure, backbone angles, contact numbers and solvent accessibility. Bioinformatics, 33(18):2842–2849, September 2017. ISSN 1367-4803. doi: 10.1093/bioinformatics/btx218.

49. Ana-Maria Fernandez-Escamilla, Frederic Rousseau, Joost Schymkowitz, and Luis Serrano. Prediction of sequence-dependent and mutational effects on the aggregation of peptides and proteins. Nature Biotechnology, 22(10):1302–1306, October 2004. ISSN 1087-0156. doi: 10.1038/nbt1012.

50. Peter Rice, Ian Longden, and Alan Bleasby. EMBOSS: The European Molecular Biology Open Software Suite. Trends in Genetics, 16(6):276–277, June 2000. ISSN 0168-9525. doi: 10.1016/S0168-9525(00)02024-2.

51. John F Peden. Analysis of Codon Usage. PhD Thesis, University of Nottingham, 1999.

52. Eric J. Ma and Arkadij Kummer. Reimplementing Unirep in JAX. bioRxiv, page 2020.05.11.088344, May 2020. doi: 10.1101/2020.05.11.088344. Publisher: Cold Spring Harbor Laboratory Section: Confirmatory Results.

53. M B B Gutierres, C B C Bonorino, and M M Rigo. ChaperISM: improved chaperone binding prediction using position-independent scoring matrices. Bioinformatics, 36(3):735–741, February 2020. ISSN 1367-4803. doi: 10.1093/bioinformatics/btz670.

54. Shifu Chen, Yanqing Zhou, Yaru Chen, and Jia Gu. fastp: an ultra-fast all-in-one FASTQ preprocessor. Bioinformatics, 34(17):i884–i890, September 2018. ISSN 1367-4803. doi: 10.1093/bioinformatics/bty560. Publisher: Oxford Academic.

55. Marcel Martin. Cutadapt removes adapter sequences from high-throughput sequencing reads. EMBnet.journal, 17(1):10–12, May 2011. ISSN 2226-6089. doi: 10.14806/ej.17.1.200. Number: 1.

56. Heng Li and Richard Durbin. Fast and accurate short read alignment with Burrows-Wheeler transform. Bioinformatics (Oxford, England), 25(14):1754–1760, July 2009. ISSN 1367-4811. doi: 10.1093/bioinformatics/btp324.

57. Heng Li, Bob Handsaker, Alec Wysoker, Tim Fennell, Jue Ruan, Nils Homer, Gabor Marth, Goncalo Abecasis, Richard Durbin, and 1000 Genome Project Data Processing Subgroup. The Sequence Alignment/Map format and SAMtools. Bioinformatics (Oxford, England), 25 (16):2078–2079, August 2009. ISSN 1367-4811. doi: 10.1093/bioinformatics/btp352.

58. Simon Anders, Paul Theodor Pyl, and Wolfgang Huber. HTSeq—a Python framework to work with high-throughput sequencing data. Bioinformatics, 31(2):166–169, January 2015. ISSN 1367-4803. doi: 10.1093/bioinformatics/btu638.

