## supplementary_information for "Experimental characterisation of *de novo* proteins and their unevolved random-sequence counterparts"

Accompanying the manuscript: '**Experimental characterisation of *de novo* proteins and their unevolved random-sequence counterparts**'

Brennen Heames[1], Filip Buchel[3,4], Margaux Aubel[1], Vyacheslav Tretyachenko[3], Andreas Lange[1], Erich Bornberg-Bauer[1,2*], Klara Hlouchova[3,5*]

[1]Institute for Evolution and Biodiversity, University of Münster, Germany
[2]Department of Protein Evolution, MPI for Developmental Biology, Tübingen, Germany
[3]Department of Cell Biology, Charles University, BIOCEV, Prague, Czech Republic
[4]Department of Biochemistry, Charles University, Prague, Czech Republic
[5]Institute of Organic Chemistry and Biochemistry, Czech Academy of Sciences, Prague, Czech Republic

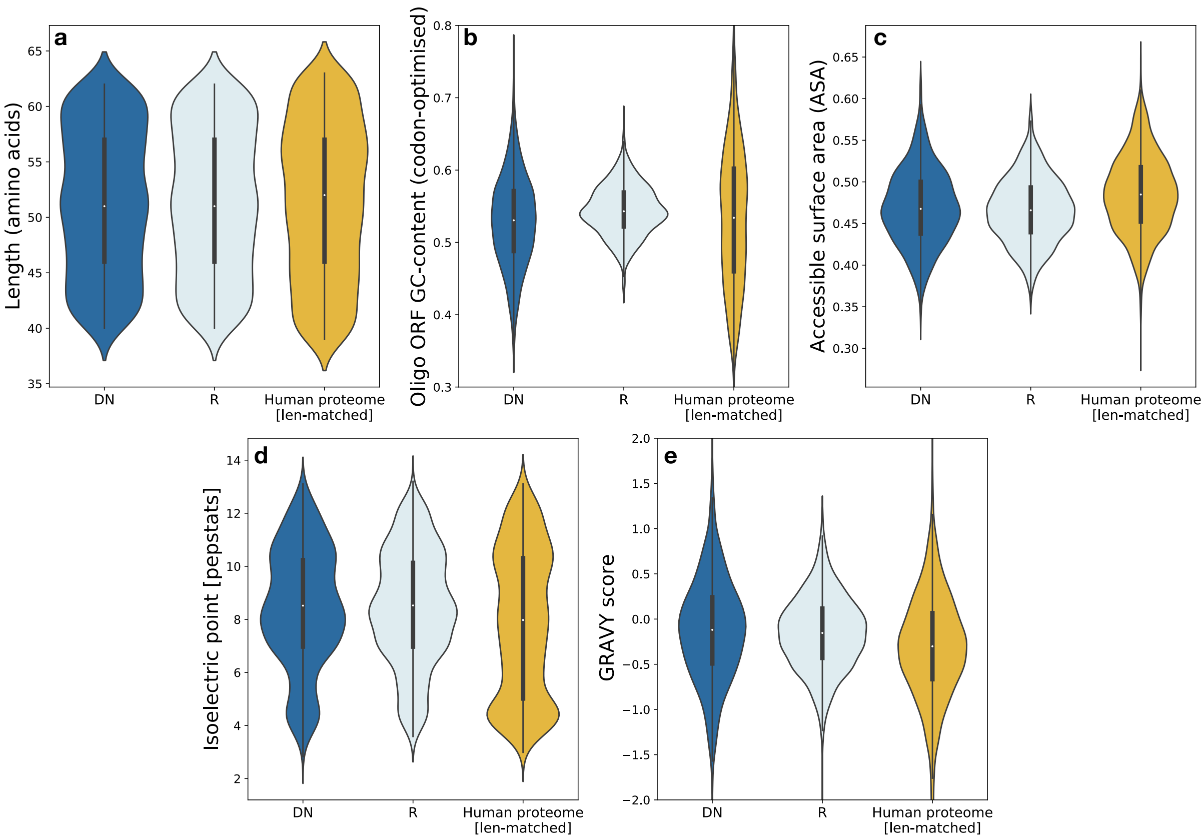


Figure S1: **Predicted sequence properties for a synthesised library of de novo and synthetic random sequences.** Distribution of each property shown for library DN (n=1800; dark blue), library R (n=1800; pale blue), and a random, length-distribution matched sample of proteins from the human proteome (n=3600; yellow).

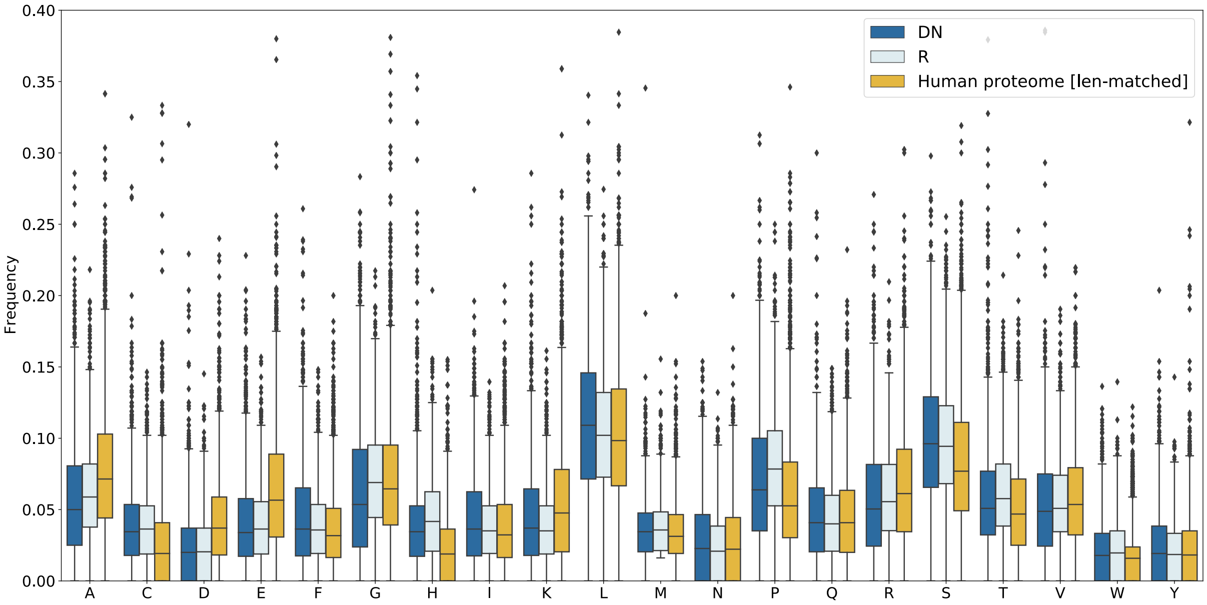


Figure S2: **Amino acid composition of libraries DN and R compared to a set of conserved human proteins.** A synthetic random protein (R) library was designed to have closely matched length distribution and amino acid composition as the library of putative *de novo* proteins (DN). A length-matched random sample of the annotated human proteome (distinct from the putative human *de novo* proteins in library DN) is shown as a reference.

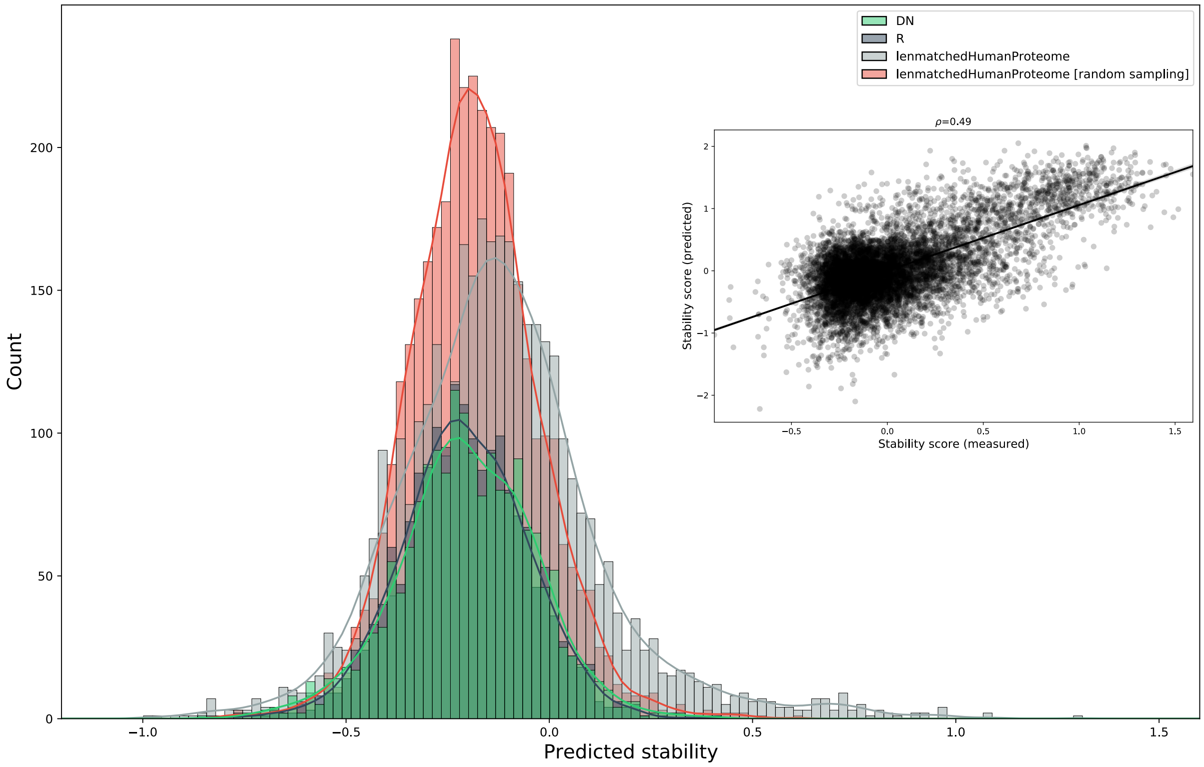


Figure S3: **Prediction of library stability using a learned protein embedding.** The UniRep embedding was used to train a model based on proteolysis-derived stability scores. Using this model, library R, with length and amino acid frequency distributions identical to library DN, is predicted to have a similar stability distribution. Predictions for a length-matched subset of the annotated human proteome (grey) and for a randomised equivalent (red) are shown for comparison. **Inset:** Correlation of predicted and measured stability scores on holdout set after training on experimental dataset of ca. 50,000 de novo designed proteins assayed by Rocklin *et al.* (2020).

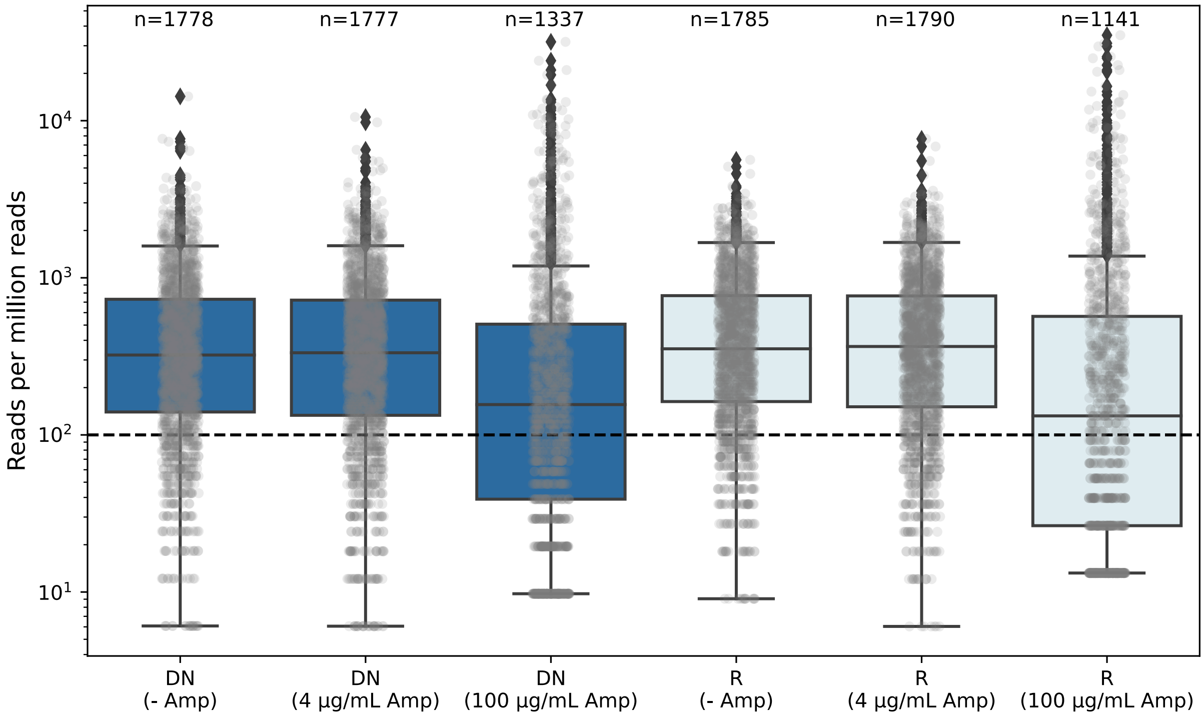


Figure S4: **Twin-arginine export quality assay: NGS read count distributions.** Read counts normalised for sequencing depth across demultiplexed sample subsets. The number of unique variants with one or more mapped reads is indicated above each bar (of a possible 1800 variants). Based on read count distributions, a threshold of 100 reads-per-million (dashed line) was used to remove those variants present at only low levels in the input library. Fractions of each library represented above this threshold are show in Fig. 3 and Fig. S6. The distribution of read counts was highly similar for libraries DN and R following initial PCR amplification and sub-cloning (i.e. comparing input libraries, '-Amp'), and represented the designed libraries with >1770/1800 variants present after drop-out.

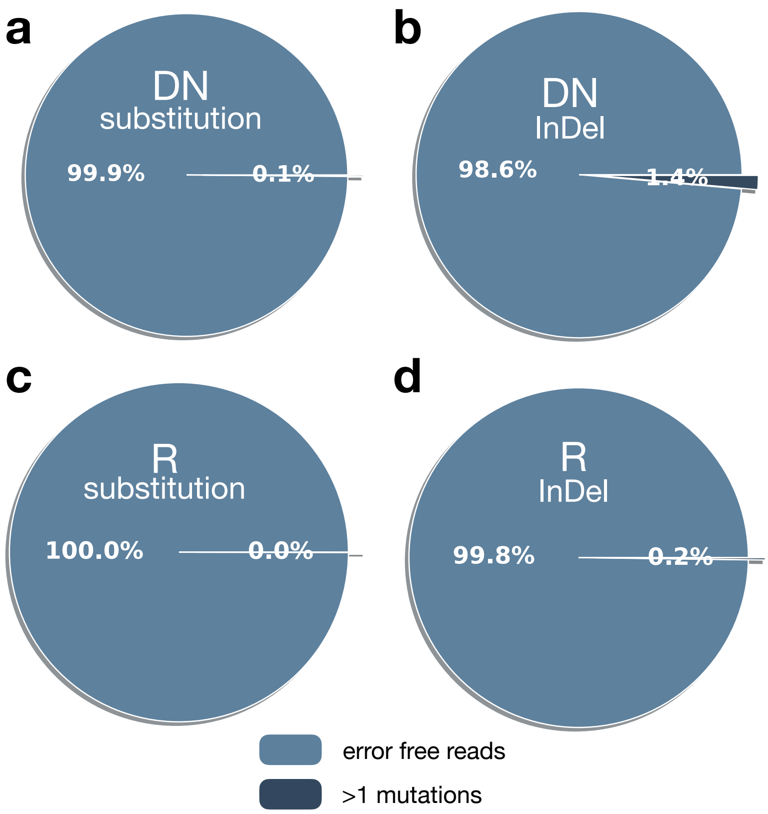


Figure S5: **Library quality following synthesis**. Next-generation sequencing (NGS) was used to assess the diversity and quality of each library. Error rates calculated independently for substitutions (**a,c**) and InDels (**b,d**) show that the majority of reads were error free following synthesis and sub-cloning of both libraries DN and R.

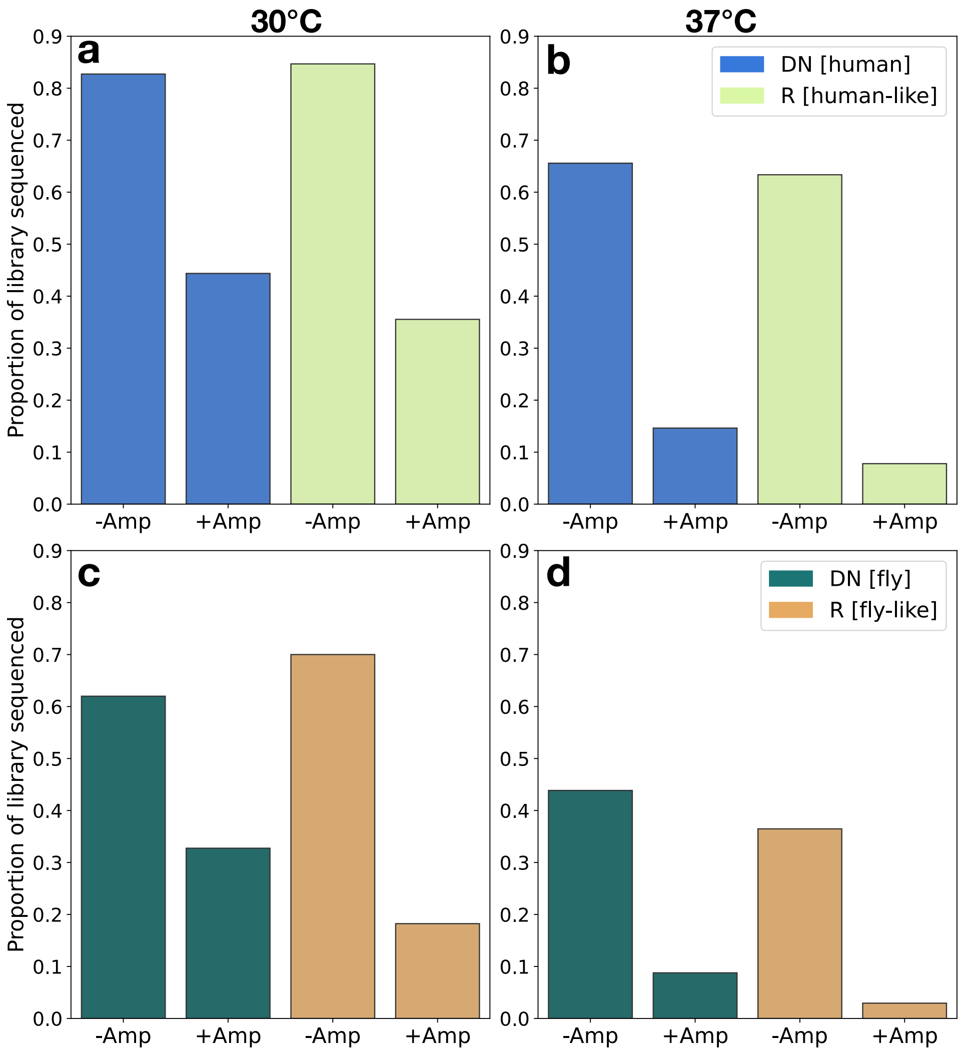


Figure S6: **Twin-arginine assay NGS survival quantification.** Proportion of library identified by NGS following plating with or without ampicillin. Bars are normalised by size of each library subpool (1624 human and human-like proteins; 176 fly and fly-like proteins) **a)** & **b)**: human *de novo* and human-like random subsets **c)** & **d)**: fly *de novo* and fly-like random subsets.

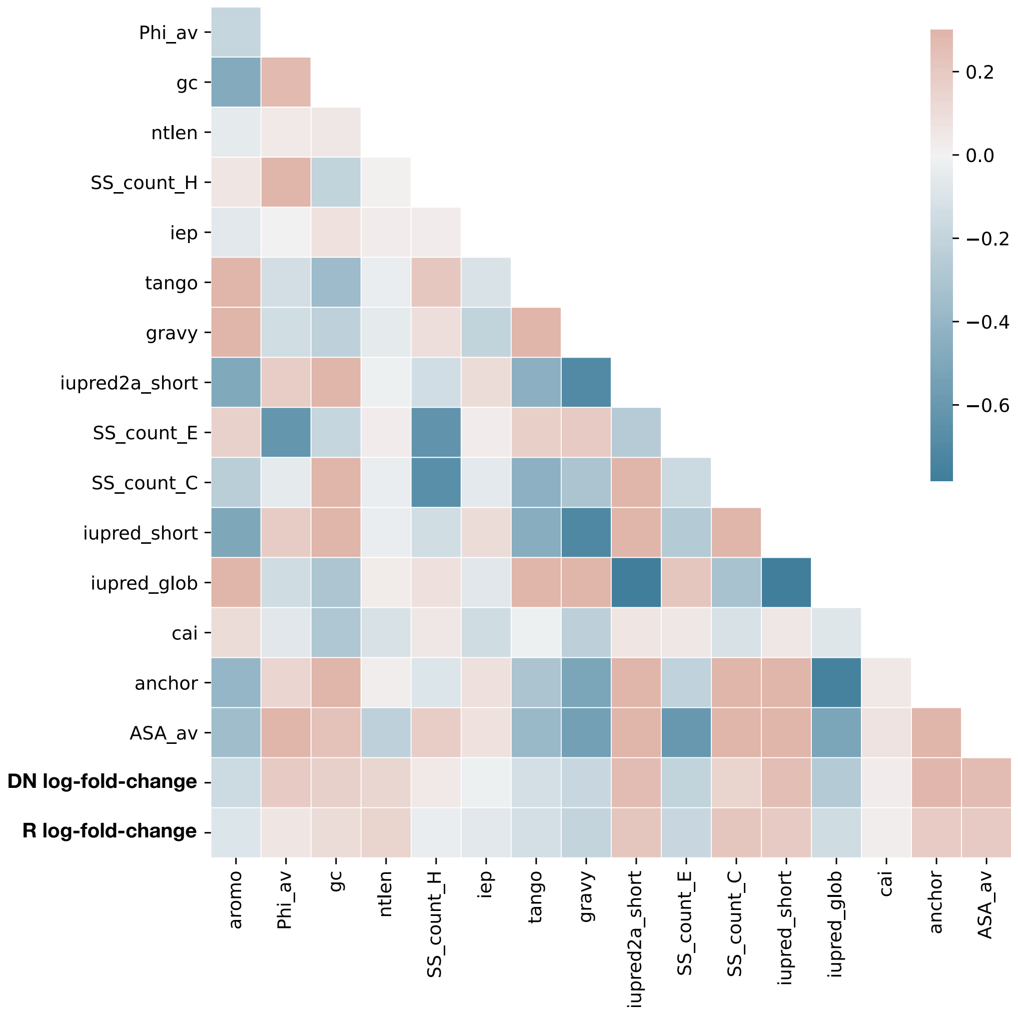


Figure S7: **Correlation matrix of predicted sequence features.** Pearson correlation coefficient between pairs of predicted protein properties indicated by colour. Features are also shown correlated with Twin-arginine assay log2-fold-change (i.e. enrichment following selection on ampicillin): 'DN log-fold-change' & 'R log-fold-change'.

| **Primer** | **Sequence** |
|---|---|
| DN_F | CTATGA gaattc GAGTCCCATGTACATATG |
| DN_R | TCATAG ggatcc TGACGACCCCTGCTCGAG |
| R_F | CTATGA gaattc TGGCTCCATACACATATG |
| R_R | TCATAG ggatcc CAGAGGAACCCGCTCGAG |
| pSAL_NSG_F_01 | CGTACTAG gactgcgcatatggaattc |
| pSAL_NSG_F_02 | AGGCAGAA gactgcgcatatggaattc |
| pSAL_NSG_F_03 | TCCTGAGC gactgcgcatatggaattc |
| pSAL_NSG_F_04 | TAGGCATG gactgcgcatatggaattc |
| pSAL_NSG_R_01 | TATCCTCT gtttctgggtgggatcc |
| pSAL_NSG_R_02 | CTCTCTAT gtttctgggtgggatcc |

Table S1: **Primers used in this study.** DN and R subpool primers (top) were used to amplify each subpool from the total oligonucleotide pool and introduce EcoRI and BamHI restriction sites prior to sub-cloning. 'NGS' primers, used for amplicon preparation prior to Next-Generation Sequencing, encode 8-bp barcodes to allow pooling of multiple conditions in a single sample.
